# Glutamatergic neurons in the infralimbic cortex for motivational response of pain relief induced by electroacupuncture

**DOI:** 10.1101/2023.03.02.530902

**Authors:** Hui Liu, Meiyu Chen, Jiaqi Lu, Chuan Qin, Can Wang, Sheng Liu

## Abstract

Peripheral neuromodulation by electroacupuncture (EA) is a promising tool for both experimental and clinical applications. However, whether EA signals reflect a multidimensional composite and evoke affective and motivational processes remains largely elusive. Here, we demonstrated that EA at ST.36 acupuncture point considerably attenuated pain hypersensitivity at 24h and 48h postincision. In conditioned place preference (CPP) model, one chamber becomes associated with EA through three-day repeated pairings, whereas the other chamber is associated with no EA stimulation. EA stimulation resulted in strong preference for the chamber paired with EA in incisional injury (INP) rats. In contrast, EA at non-acupuncture points in INP rats did not relief pain and produce CPP. Notably, EA with the context in sham-operated animals did not induce CPP. Next, we identified neurons activation in brain associated with affective and motivational aspects of pain after EA stimulation using immediate early gene c-Fos expression in SNI rats. EA stimulation increased c-Fos positive neurons in the IL, but not cingulate (Cg1) and prelimbic (PL) subregion of the mPFC. Sham EA did not increase c-Fos expression in the IL in spared nerve injury (SNI) rats. Using reversible inactivation of IL in rats, inactivation of the IL significantly abolished CPP of pain relief induced by EA. Optogenetic activation of IL glutamatergic neurons mimicked EA-induced analgesia and CPP behaviors, and inhibition of glutamatergic neurons in the IL reversed the effects of EA. The study directly demonstrates a novel and important role for glutamatergic neurons in the infralimbic cortex in acupuncture-induced motivational response of pain relief and provides a new perspective for investigating acupuncture analgesia.

## Introduction

Almost 5000 years ago, ancient people have learned to stimulate peripherally some sites of their body to relief various diseases or conditions [1]. Peripheral neuromodulation has become worldwide in its practice after several decades of rapid development and brought new insights into the diagnosis and treatment of diseases [2]. A growing body of evidence supports the use of acupuncture or electroacupuncture (EA), one of popular peripheral neuromodulation techniques targeting a stimulus to specific points in the body, as an effective treatment for acute and chronic pain, including headaches, neck pain, osteoarthritis pain and others [3-5]. Its analgesic mechanisms were explained partly by the gate control theory, and involved somatosensory autonomic reflexes, peripheral and central opioid, dopamine, noradrenaline, and serotonin systems [6,7]. These experimental efforts of EA analgesia have been largely focused on the perception of pain [8]. In addition, the various observations that have been made are not sufficient to permit a unified theory regarding the analgesic effect of EA.

The current conceptualizations of pain in humans are predominantly multidimensional, mainly including the perception of the noxious stimulus and the affective-motivational features of pain. Experimental and clinical studies show robust reciprocal interactions between pain perception and affect, in which negative affects facilitate pain, while positive ones inhibit it [9]. Importantly, chronic pain can induced maladaptive motivational states, which lead to opioid overdose and much neuropsychiatric comorbidity [2]. Affect and motivation are thought to exert substantial influence on the subsequent pain relief. Furthermore, affective and motivational responses drive patients to seek treatment which can cause them to alter their lifestyle to avoid pain [10]. Greater emotion variability is associated with lower life satisfaction, worse psychosocial functioning, and greater depression and anxiety [23,24]. Notably, recent evidence supports peripheral neuromodulation with acupuncture or EA is more than a tool for treatment of pain perception. Beneficial effects of acupuncture or EA in psychological conditions, including anxiety, depression, and cognition, caused by chronic pain were observed in animal and human studies [11,12]. Unveiling whether and how the peripheral neuromodulation with EA integrates pain affective-motivational state is of particular interest in the context of treating pain, and would shed new light on analgesia induced by peripheral neuromodulation in different domains.

Mounting evidences suggest that EA stimulation induces widespread responses in many brain areas (such as the prefrontal cortex, the limbic system, and subcortical gray structures), which may provide important central circuitry basis for integrating pain emotion and shape behavior by peripheral stimulation with EA [13,14]. However, the specific findings of these studies have not been entirely consistent with one another, and it is not clear whether the observed alterations are a cause or a consequence of pain [3]. The medial prefrontal cortex (mPFC) may be an idea candidate for tow-down control of EA analgesia, given its role in processing complex stimulus-response information and the modulation of sensory and affective components of pain [15]. It is shown that the mPFC is deactivated in chronic pain and the activation of mPFC gave rise to analgesic effects and provided important control for the aversive response to transient noxious stimulations [16-18]. Some studies suggest that peripheral neuromodulation with EA involves mPFC modulation [19]. For example, infralimbic medial prefrontal cortex (IL) alters EA effect in neuropathic chronic pain rats [20]. Work from our lab shows that acupuncture can modulate the activity of IL neurons [21,22]. However, the precise cell-type specific organization and the function of cortex circuits mediating EA-induced analgesia remain obscure. Here, we specified the glutamatergic neurons in the infralimbic cortex involved in EA analgesia, and illustrated an example in which affective-motivational dimension of pain is directly influenced through top-down control of the peripheral neuromodulation with EA.

## Materials and Methods

### Animals

Adult male Sprague-Dawley rats (250–350 g; Slack) were purchased from the Shanghai Laboratory Animal Center at Chinese Academy of Science (Shanghai, China). Male C57BL/6N (6-8 weeks old), purchased from SLAC laboratory (Shanghai), and CamKIIα-Cre (6-8 weeks old), a gift from Zhang Yuqiu’s lab at Fudan University, were used for experiments. All animals were raised in stable conditions maintained at a controlled temperature of 23±3 °C and air humidity of 40–70% under 12-hr light/dark cycle (lights on from 7:00 A.M. to 7:00 P.M.) with ad libitum food and water. Behavioral tests should be performed at the time of the light and the animals were allowed to adapt to the environment for 1 week before the experiment was performed. All experiments were approved by the Institutional Animal Care and Use Committee and the Animal Ethics Committee (SZY 201710008) of Shanghai University of Traditional Chinese Medicine. The experiment was conducted in the Laboratory of Experimental Acupuncture, Shanghai University of Traditional Chinese Medicine, in accordance with local guidelines for animal welfare.

### Animal models of Incisional Injury Pain

Incision injury of the skin plus deep tissue, including fascia and underlying muscle, was done as described by Woo et al. [9]. We anesthetized rats with 2% (vol/vol) isoflurane, and made a 1-cm longitudinal incision through the skin of the left hind paw. We elevated the plantaris muscle and incised longitudinally. We stitched the cut skin with two 5–0 nylon sutures and the wound site treated with neomycin. We anesthetized sham animals and cleaned the left hind paw, but made no incision.

### Animal models of Spared Nerve Injury (SNI)

As described in the previous studies [10,11], We anesthetized the rats with isoflurane (5%). We made an incision proximal to the lateral side of the right knee to separate the biceps femoris and to expose the sciatic nerve and its associated branches. We separated the biceps femoris. We exposed the sciatic nerve and its branches. Then, we used a silk suture to ligate the common peroneal and tibial nerve branches. We removed about 1 mm of the nerve and kept the sural nerve intact. We performed the operation by following the same procedure in the Sham SNI group, but we did not cut-off or ligate the sciatic nerve, peroneal nerve, and tibial nerve branches.

### EA treatment

We restrained rats gently. Rats were submitted to EA with two stainless steel needles (0.16 mm diameter, 13 mm length) inserted into bilaterally ST36, SP6 or BL23 at a depth of 5mm. Acupuncture point ST36 is located at near the knee joint, between the muscle anterior tibialis and muscle extensor digitorum longus. We used a stimilator (Model G-6805-2, Shanghai Medical Electronic Apparatus, China) via the needles. Stimulation frequency was 2 Hz. The intensity of the stimulation was 3 mA was increased stepwise from 0.5 to 1.5 mA, with each step lasting for 10 min. As the sham control, the needles inserted into ST36 but electric stimulation was not conducted.

For mice, acupuncture needles of 0.16 mm in diameter were inserted at a depth of 2-3 mm into bilateral Zusanli (ST36). The parameters were set as follows: 2 Hz frequency, 2 mA intensity for a total of 30 min. For sham EA treatment, mice were inserted with acupuncture needles but without electrical stimulation.

### Nociceptive behavior

For examining thermal pain, we used an IITC Model 390 Paw Stimulator Analgesia Meter (IITC/Life Science Instruments, United States) to measured paw withdrawal latency (PWL). We placed the rats or mice in an inverted plastic cage. After the rats or mice were accommodated for 30 min, we used a radiant heat to warm the hind paw of rats or mice until they lifted their paw. We adjusted the radiant heat intensity to induce the response around 12–13 s in a normal rat or mice. Thereafter, we analyzed the PWL scores by calculating the time from onset of radiant heat application to the paw withdrawal. The test was repeated three times independently in each animal to calculate the mean values of the PWL scores. For testing mechanical sensitivity, we placed mice in a plexiglass box with a grid at the bottom and adjusted for at least 30 minutes before allodynia measurement. We applied a series of Von Frey filaments (0.16 g as a starting dose) to vertically stimulate the hind paw of the mice according to the method of “up and down” for measuring mechanical allodynia.

### Conditional Place Preference (CPP)

We used a standard two chamber balanced design to test the affective and motivational responses after pain relief induced by EA. On the preconditioning day, we allowed animals to travel the chambers freely for 15 min, and recorded the amount of time spent in each chamber. We analyzed data using behavior analysis system (Shanghai Jiliang Software Technology Co., Ltd). Next, two conditioning sessions (for EA and gentle handling) were performed for three consecutive days. We treated animals with either EA or gentle handling for 30 min each in the morning and afternoon. We immediately placed the rats into the pairing chamber and confined them to the chamber for 30 min after administration of 20 min EA or gentle handling. For every conditioned animal, We carried out EA and gentle handling alternatively in the morning and afternoon sessions. The morning and afternoon EA/gentle handling were conducted at least 4 h apart. We conducted a preference test 3 days after the conditioning. Then, we placed animals in CPP apparatus with a door opened in the middle for 15 min to allow access to the apparatus’s whole parts in this phase. We analyzed the preference scores by calculating the amount of time the animals spent in the EA reward paired chamber minus the time they spent in the gentle handling chamber. Sham EA group was conditioned with Sham EA treatment.

### Immunofluorescence staining

We anesthetized rats using pentobarbital sodium (100 mg/kg, ip), sequentially perfused transcardially with saline (200 mL) and 4% paraformaldehyde (PFA) in 0.1 mol/L phosphate buffer (PB) (250 mL). We collected brains from each animal. The animals’ brains were fixed overnight in 4% PFA, and transferred to 30% sucrose for 5–7 days until the tissue was saturated and sank to the bottom of the solution. We cut samples to yield a series of 30 μm-thick coronal sections with a chilled (−25°C) cryostat instrument.

We washed the sections three times with PBS, blocked them for 2 h with 1% BSA at 4°C, and probed for 48 h with the primary antibodies, including mouse anti-c-fos primary antibody (ab208942,1:1000, Abcam) and VGLUT2 Polyclonal Antibody (PA5-77432,1:400, Thermo Fisher Scientific) for 48h. Following three subsequent washed with PBS, we stained sections for 2 h with primary antibodies (1:500). Then, we mounted sections on adhesive slides. We used a Leica Laser Scanning Confocal Microscope to analyze these stained sections. We identified the best standard stereotaxic plane sections of the IL as per Paxinos and Watson’s atlas [6], with numbers of positive cells per section then being assessed (20 ×). Next, we used an automatically generated 250×600 μm rectangle to denote the IL in each section. Finally, we determined numbers of positive nuclei per section for analysis.

### Surgery for viral injection into the IL

We anesthetized mice with 1% sodium pentobarbital through intraperitoneal injection with 50 mg/kg and then fixed them on a stereotaxic apparatus. We exposed the skull of mice through a median incision. Then, we used the microinjection glass pipette to locate the target brain area according to the mouse atlas of Paxinos and Watson. In order to manipulate glutamatergic neurons in IL, we injected rAAV-Ef1 α -DIO-hChR2-(H134R) -EYFP-WPRE-pA (2.0 × 1012 vg/ml) into right IL (relative to bregma: anteroposterior (AP), 1.6 mm; mediolateral (ML), −0.25 mm; dorsoventral (DV), −2.1 mm) or rAAV-Ef1α-DIO-eNpHR3.0-EYFP-WPRE-pA (3.56×1012 vg/ml) into bilateral IL (relative to bregma: anteroposterior (AP), 1.6 mm; mediolateral (ML), ±0.25 mm; dorsoventral (DV),−2.1 mm) at a volume of 80 nL with 20 nl/min in CamK II α-Cre mice. We used AAV2/9-hEf1α-DIO-EYFP-WPRE-pA (0.48×1012 vg/ml) as a control.

### Optic fiber implantation (for the fiber photometry and optogenetic experiments)

Following the viral injection, we implanted an optic fiber (diameter, 200 μm; numerical aperture (NA), 0.37, Shanghai June Biotechnology Co., Ltd) into the above right IL (relative to bregma: anteroposterior (AP), 1.6 mm; mediolateral (ML), −0.25 mm; dorsoventral (DV), −1.9 mm). To optogenetically inhibit glutamatergic neurons in IL during electroacupuncture stimulation, we implanted the optic fibers bilaterally above IL at the angle of 20 degrees (relative to bregma: anteroposterior (AP), 1.6 mm; mediolateral (ML), ±1.01 mm; dorsoventral (DV), −2.03 mm) after the bilateral IL was injected with the virus. Finally, we used gel (Vetbond Adhesive, 3M) was applied to the skull surface and dental cement to cover the exposed skull and fix the optical fibers. The animals were allowed to recover from anesthesia on an electric blanket before returning to their home cage. We performed subsequent experiments three weeks after virus expression.

### Optogenetic manipulations

We connected the implantable optic fibers to a laser generator using optic fiber sleeves. We controlled the delivery of blue light (473 nm, 6mW, 10 ms per pulse, 10Hz) or yellow light (594 nm, 5-8mW, constant) with Doric system (Doric Lenses, Canada) for 30 min. We applied the same stimulus pattern in the control group. We performed optogenetic manipulations in the conditioning phase of CPP paradigm.

### Intracranial microinjection

We injected rats (300–350 g) intraperitoneally with pentobarbital sodium (50 mg/kg) and mounted rats on a stereotaxic frame. Then, we implanted bilateral guide cannulae (26 gauge, Plastics One) in the IL (+3.0 mm AP; ± 0.8 mm ML; –3.8 mm DV). We fixed the cannulae to the skull with dental cement with three steel screws. Rats were systemically treated with benzylpenicillin sodium (60,000 U) to prevent infection. We allowed rats to recover for 7 days after surgery. Five minutes prior to EA stimulation, rats were bilaterally injected with a combination of the GABAB agonist baclofen and the GABAA agonist muscimol (B/M) (0.3 and 0.03 nmol, respectively) or same volume of DMSO into the IL. We inserted injection cannulae into the guide cannulae. We attached the injectors to a microinfusion pump (RS Instruments) via PE 10 tubing. We left the injection cannulae in place for an additional 2 min.

### Statistical analysis

Statistical analysis was performed using GraphPad Prism 7 and MATLAB. We analyzed the data using unpaired t-test, paired t-test, one-way ANOVA with Bonferroni’s correction for multiple comparisons, two-way ANOVA, Wilcoxon signed rank test and Fisher’s LSD test. p < 0.05 was the cutoff of significance for these analyses.

## Results

### Peripheral neuromodulation with EA induces motivation response after pain relief in rats of incisional injury pain

We first investigated whether EA induced affective-motivational behaviors in different animal models of pain. Preclinical studies demonstrate relief of pain induced by analgesic agents produces CPP, regardless of the animal model [25]. We therefore tested whether analgesic effect of EA produced CPP. We chose 2 Hz EA bilaterally at points ST36 (a depth of 5 mm near the knee joint, between the tibialis anterior and extensor digitorum longus muscles, Fig.1B) as a result of the experience from our previous study [26,27] and some systematic review of literature [8,28,29]. EA at ST.36 acupuncture point considerably attenuated pain hypersensitivity at 24h and 48h postincision (Fig.1C). In CPP model, one chamber becomes associated with EA through three-day repeated pairings, whereas the other chamber is associated with no EA stimulation (Fig.1A). Preference scores is measured by the amount of time animals spend in the EA associated chamber minus the time it spends in the non-EA chamber, when given free access to both chambers after conditioning. EA stimulation resulted in strong preference for the chamber paired with EA in INP rats. In contrast, EA at non-acupuncture points in INP rats did not relief pain and produce CPP (Fig.1D,E). Notably, EA with the context in sham-operated animals did not induce CPP, indicating that EA is not motivational in the absence of pain initially. CPP was induced by EA only when there is pain.

**Fig.1.**
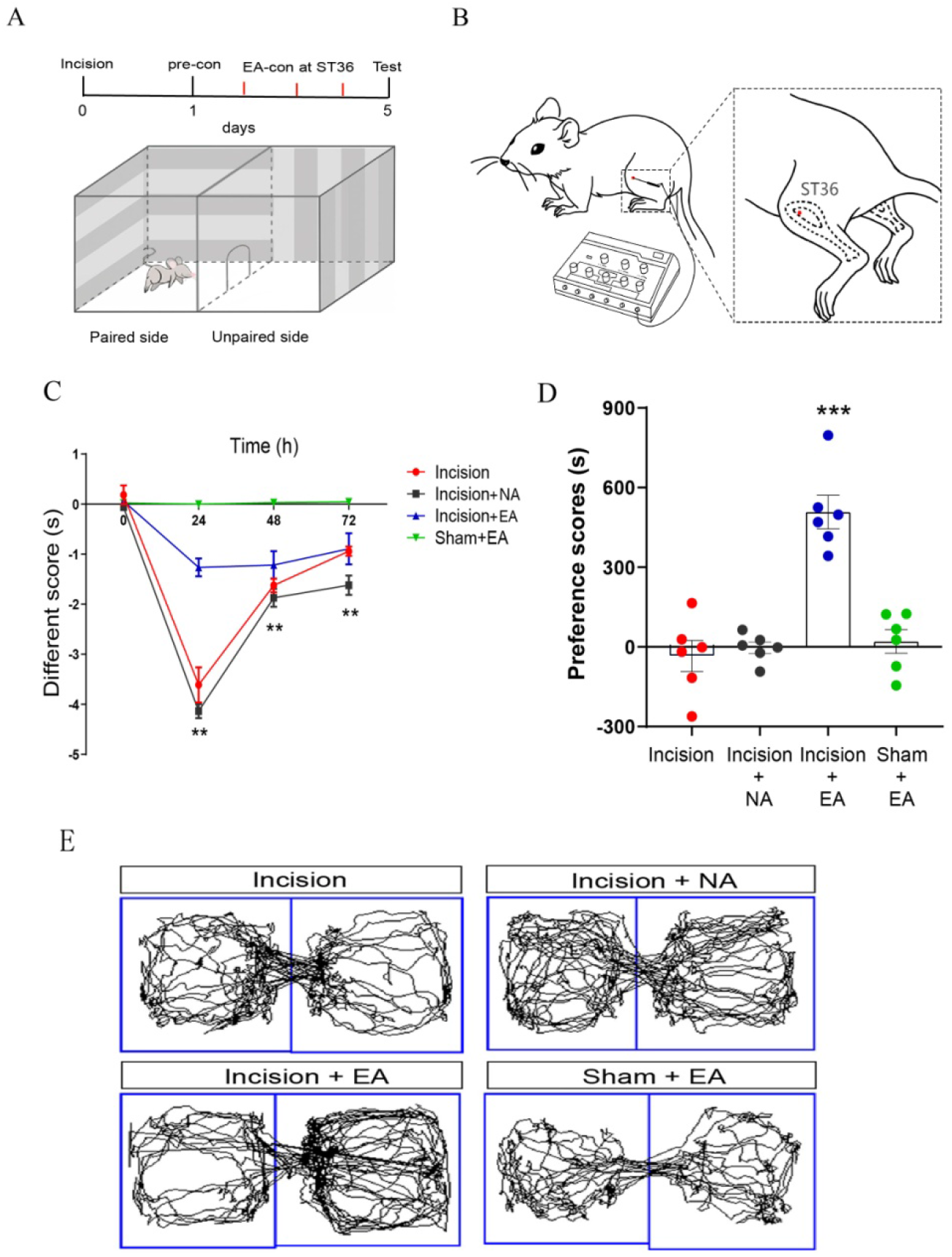
EA induced analgesia and CPP in rats of incisional injury pain. (A) Schematic showing the protocol of experiment. (b) Schematic representation of placement of EA stimulation at ST36. (C) EA attenuated pain hypersensitivity in INP rats. n=6 rats in each group. Different scores, which was calculated by subtracting the PWL of the untreated paw from the PWL of the injured paw. (two-way ANOVA for repeated measures: time×group: F9,60 = 26.9, p < 0.0001). **p < 0.01 versus Incision, EA at non-acupoints (Incision+NA), and EA in sham operated animals (Sham+EA). (D) Preference scores after CPP testing among the different groups expressed as the time spent in the EA-paired side minus the time spent on the non-paired side on the test day. ***p < 0.001 vs. Incision, Incision+ NA, and Sham+EA. (E) Real-time movement traces among the groups during CPP test.

In C57 mice with spared nerve injury (SNI), we found that the thresholds of mechanical allodynia and thermal hyperalgesia were significantly decreased after SNI modelling, while EA stimulation could relieve the mechanical allodynia and thermal hyperalgesia (Fig2 C,D). The accumulative analgesia of EA was maintained for at least 45 days after SNI. Animals were given EA pairing CPP training on 7 to 11 days after SNI modeling (Fig.S2 A,B). EA produced a significant preference for the chamber paired with EA that was not found in a group of animals given a sham EA (Fig.2F-G), suggesting that EA also induced CPP in animals with chronic neuropathic pain.

**Fig.2.**
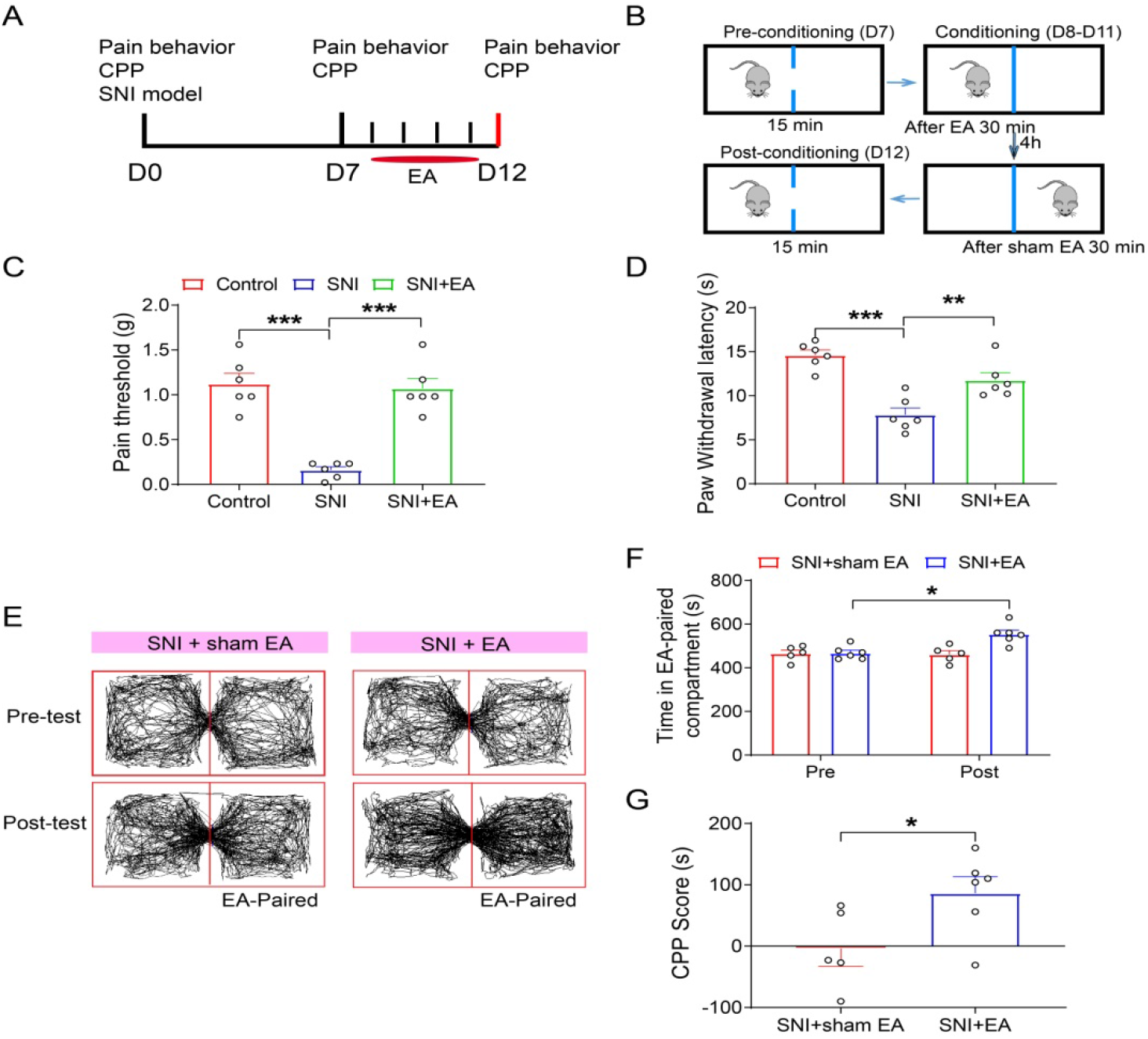
EA induces analgesia and CPP behavior in SNI mice. (A) Schematic showing the protocol of experiment. (B) Procedure to elicit and test EA induced-CPP behavior. (C-D) The thresholds of mechanical allodynia (C) and thermal hyperalgesia (D) were decreased in SNI mice. The thresholds of mechanical allodynia and thermal hyperalgesia were increased after EA stimulation. n=6 mice in each group. (E) The trajectory of SNI mice in a CPP apparatus during pre-test phase and post-test phase. (F) After four consecutive days of EA stimulation, the residence time of EA-conditioned chamber were increased in SNI mice. The residence time of SNI mice in CPP apparatus was not affected by sham EA. (G) The CPP score was increased after EA not sham EA stimulation in SNI mice. n=6 mice for EA and n=5 for sham EA. *p < 0.05, **p < 0.01, ***p < 0.001. All data were presented as mean ± s.e.m. One-way ANOVA followed by Bonferroni’s test (C and D), Paired Student’s t test (F) and Unpaired Student’s t test (G) were used for statistical analysis.

### EA activates glutamatergic neurons in the infralimbic (IL) cortex in rats of pain

Pyramidal neurons in the mPFC produce glutamate as a neurotransmitter and glutamatergic neurons are the major projection neurons in the IL. We hypothesized that glutamatergic projection neurons expressing vesicular glutamate transporter 2 (vGlut2) in the IL may play a central role in EA modulation effect on pain processing. To confirm this, we performed c-Fos/vGlut2 double labeling to examine whether glutamatergic projection neurons were activated in SNI rats following EA stimulation (Fig.3A). EA stimulation on day 14 after SNI modeling increased the numbers of c-Fos-positive vGlut2 labeling neurons in the IL (Fig.3B-D). EA did not change the percentages of vGlut2 staining neurons that were c-Fos positive in the IL in sham operated rats. These data demonstrate that EA stimulation increased c-Fos expression preferentially in IL glutamatergic neurons in rats with pain.

**Fig.3.**
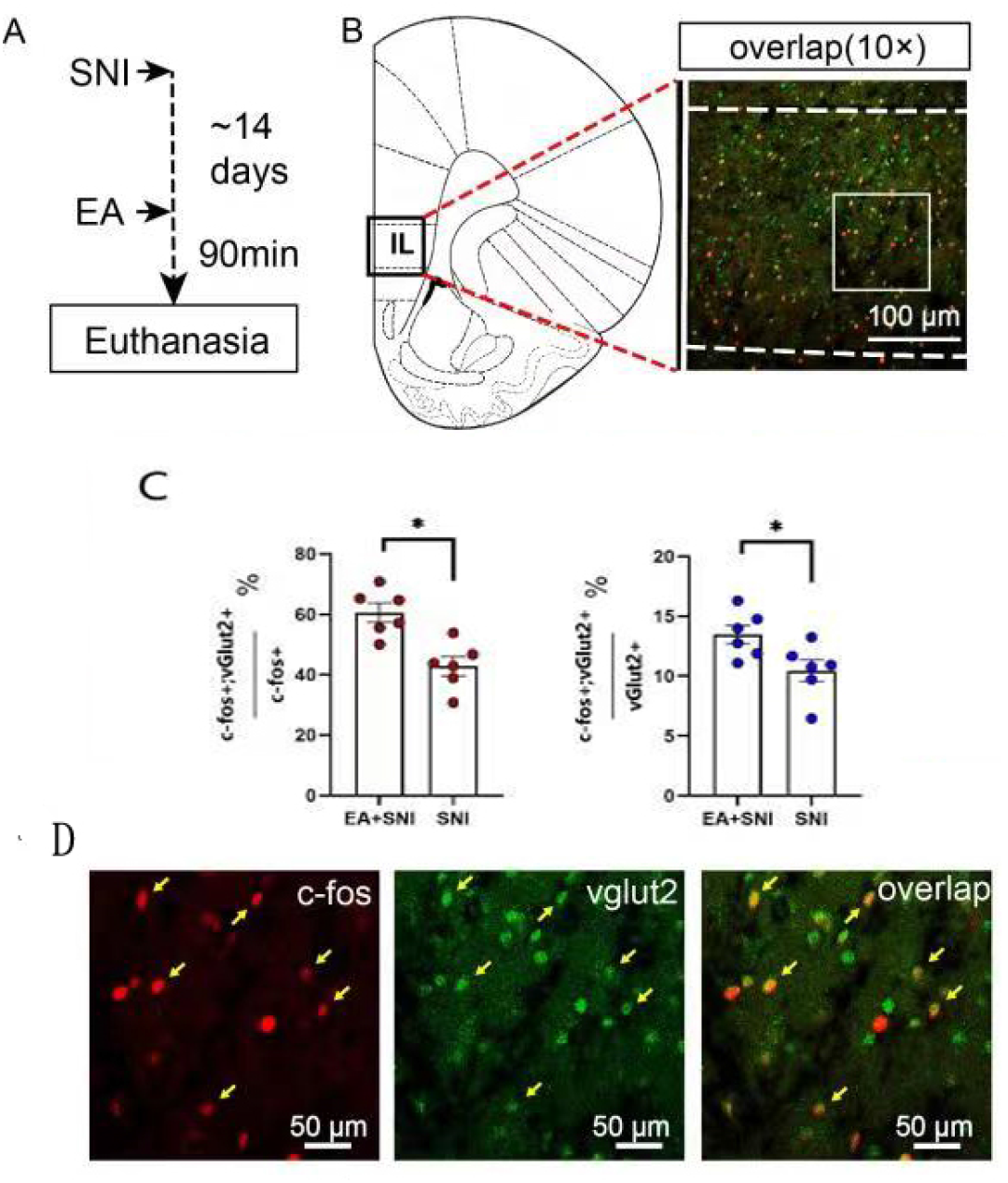
EA activates glutamatergic neurons in the IL in rats of pain. (A) Experimental strategy. (B) Representative photomicrograph showing that EA induced increase in the numbers of c-Fos-positive vGlut2 labeling neurons in the IL. Top: scale bar, 100 μm. Bottom: magnified image shows the morphology of labelled neurons in the boxed area. (C)-(D) Quantification of double-labeled vglu2/c-Fos neurons in the IL. n=6 rats in each group; P=0.002 (C); P=0.033(D). Data were expressed as mean ± s.e.m and analyzed by unpaired t-test.

### Optogenetic activation of glutamatergic neurons in the IL mimics the effect of EA in SNI animal model

Above data are correlational but do not determine whether increased neuronal activity is a cause or a consequence of EA effects. To further determine the effects of activation of IL glutamatergic neurons in the pain and affective-motivational behaviors, a cre-dependent rAAV expressing channelrhodopsin-2 (ChR2) fused with enhanced yellow fluorescent protein (rAAV-DIO-ChR2-EYFP) was injected in CamK II α-Cre mice and an optical fiber was implanted at the injection site to manipulate the glutamatergic neurons (Fig.4A,B). We found approximately 80% of ChR2 expression cells overlapped with anti-Camk II immunostaining. The mice were stimulated by 473 nm blue light to match one chamber of the CPP apparatus. Administration of blue light in the IL increased thresholds of mechanical allodynia and thermal hyperalgesia in SNI mice (Fig.4C,D), suggesting that the excitation of IL glutamatergic neurons could produce analgesic effects.

**Fig.4.**
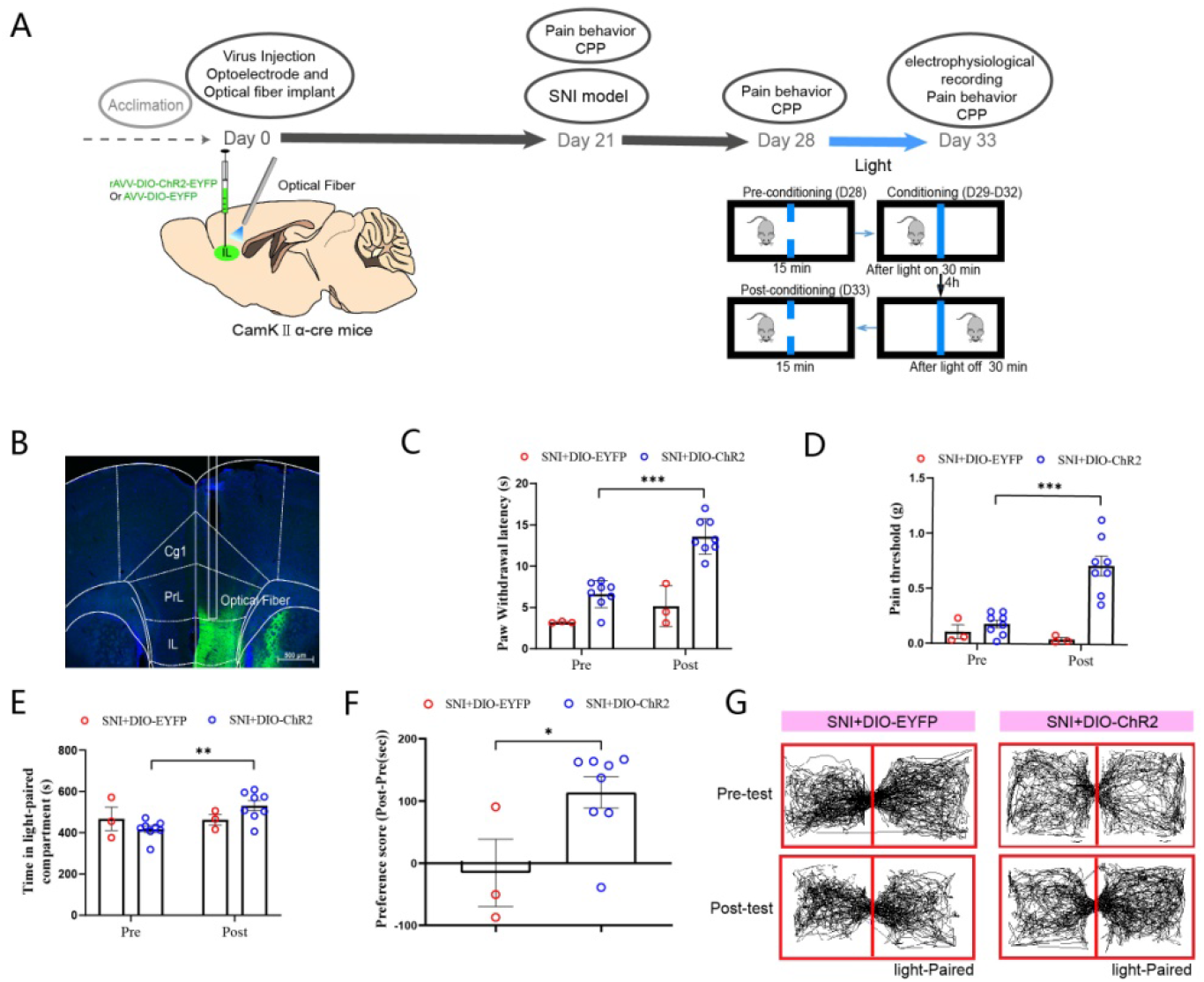
Optogenetic activation of glutamatergic neurons in the IL mimics the effect of EA in SNI animal model. (A) Schematic diagram showing the timeline of experiment and injection of AAV-EYFP or rAAV-ChR2-EYFP into the right IL and embedding of optical fiber in CamK II α-Cremice. (B) Representative images of showing viral injection site and fiber embedding site in CamK II α-Cre mice. Bar = 500 μm. (C)-(D) The thresholds of mechanical allodynia and thermal hyperalgesia were increased after blue light stimulation. (C) Paw withdrawal latency (s) before and after light stimulation; (D) Pain threshold (g) before and after light stimulation. n=3 mice for SNI+DIO-EYFP. n=8 mice for SNI+DIO-ChR2. (E)-(F) After four consecutive days of light stimulation, the residence time of light-conditioned chamber were increased in SNI model mice. (E) Time in light-paired compartment (s) before and after light stimulation; (F) The CPP score was increased after light stimulation in SNI mice. n=3 mice for SNI+DIO-EYFP. n=8 mice for SNI+DIO-ChR2. (G) The trajectory of SNI mice in a CPP apparatus during pre-test phase and post-test phase. Two-way analyses of variance (ANOVAs) followed by Fisher’s LSD test (C, D and E) and Unpaired Student’s t test (G) were used for statistical analysis.

Interestingly, delivery of blue light in the IL resulted in robust preference for the chamber paired with light stimulation in CamK II α-Cre mice with SNI (Fig.4E-G). These results indicated that the effect of IL glutamatergic neurons optogenetic activation on hyperalgesia and CPP was similar to that of EA.

### IL inactivation reverses the effects of EA in SNI mice

We reasoned that if activation of these neurons mimics the effects of EA-induced peripheral neuromodulation, then inhibition of these cells should reverse EA-induced analgesia and CPP. To test this, we first determined a causal role of IL inactivation in EA-induced CPP using a mixture of muscimol+baclofen (GABAa+GABAb agonists) to non-selectively inactivate the majority of IL neurons 5–10 min prior to EA treatment in SNI rats (Fig.5A). The rats were divided randomly into two groups (n =6 per group): EA+B/M (combination of the GABAB agonist baclofen and the GABAA agonist muscimol) and EA+Sal (saline as a control) group. As shown in Fig.5, there were significant differences between groups in preference scores. EA+B/M group spent significantly less time in EA paired compartment on the postconditioning test day (417.2 6± 34.2 s). By contrast, the same procedure did result in the production of CPP in EA+saline rats (355.8 6 ±29.9 s). It is clear that inactivation of the mPFC abolished CPP of pain relief induced by EA.

**Fig.5.**
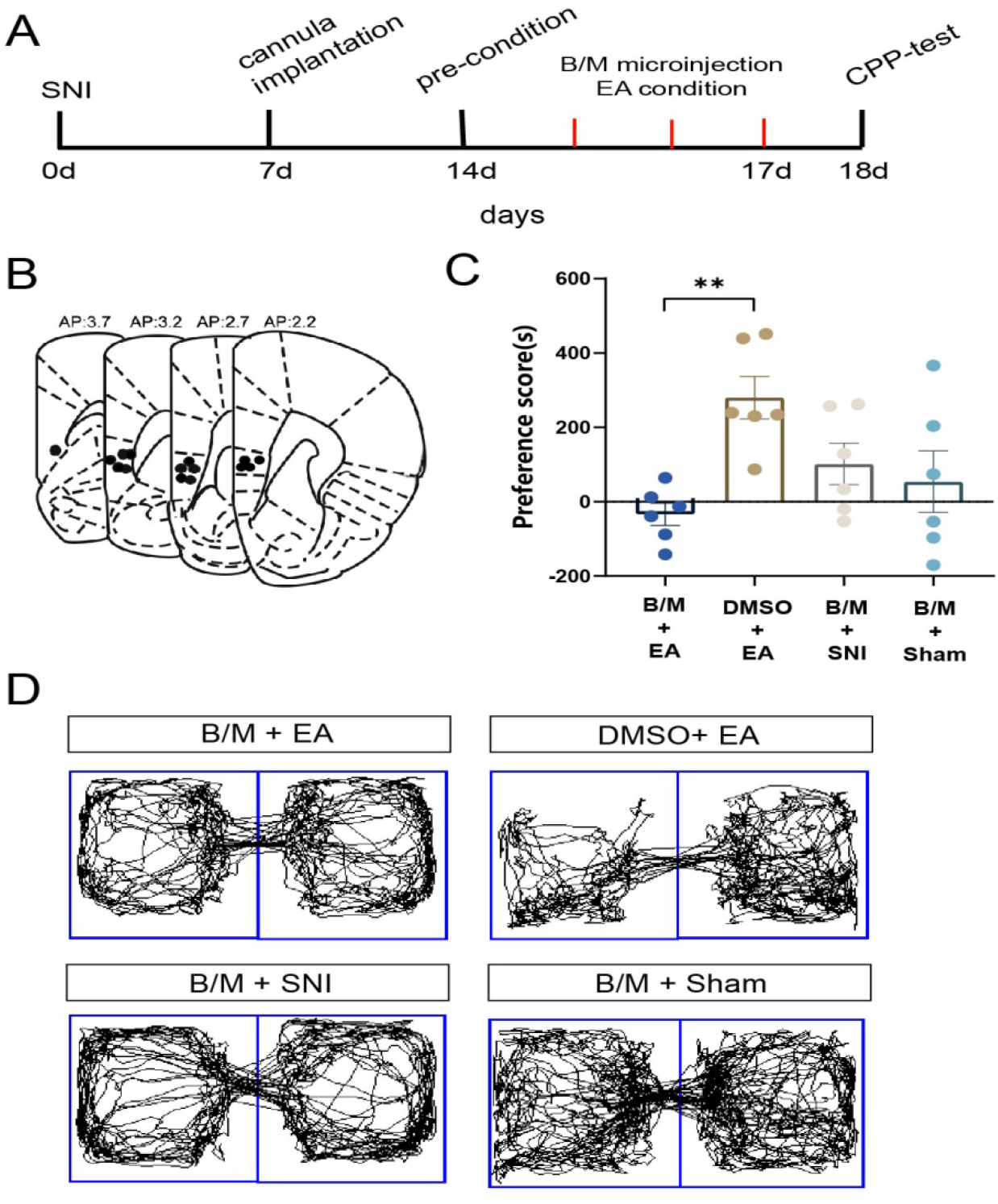
Transient IL inactivation disrupts the effects of EA on CPP behaviors in SNI model rats. (A) A timeline of SNI modeling, EA treatment, stereotactic injection, and behavioral testing protocols. (B)Microinfusion cannula placements. The numbers indicate distance from Bregma in millimeters. (C)The preference scores after CPP testing among the various groups. n=6 in each group. Data are expressed as the mean ± s.e.m. One-way analysis of variance (ANOVA) followed by Fisher’s LSD test were used for statistical analysis. **p < 0.01. (D) Real-time movement traces among the different groups during CPP test: B/M + EA group (left top); DMSO + EA group (right top); B/M + SNI group (left bottom); B/M + Sham group.

### Optogenetic inhibition of glutamatergic neurons in IL reverses the effects of EA in SNI mice

We test whether optogenetic inhibition of glutamatergic neurons in IL reverses the effects of EA in SNI mice. A cre-dependent rAAV expressing eNpHR fused with enhanced yellow fluorescent protein (rAAV-DIO-eNpHR-EYFP) was injected in CamK II α-Cre mice and an optical fiber was implanted at the injection site to inhibit the glutamatergic neurons by using 589 nm yellow light stimulation. We found approximately 80% of eNpHR expression cells overlapped with anti-Camk II immunostaining. Yellow light (594 nm, 5-8mW, constant) was delivered into the IL while SNI mice were received EA stimulation. We observed that the increased thresholds of mechanical allodynia and thermal hyperalgesia induced by EA were inhibited while yellow light was delivered simultaneously (Fig.6), indicating that inhibition of IL glutamatergic neurons reversed the analgesic effects of EA. Furthermore, EA-induced CPP (increased CPP scores) were ablated by delivery of yellow light (Fig.6E-H) in the IL. These results clearly demonstrate that glutamatergic neurons in IL are necessary for EA-induced pain relief and CPP behaviors.

**Fig.6.**
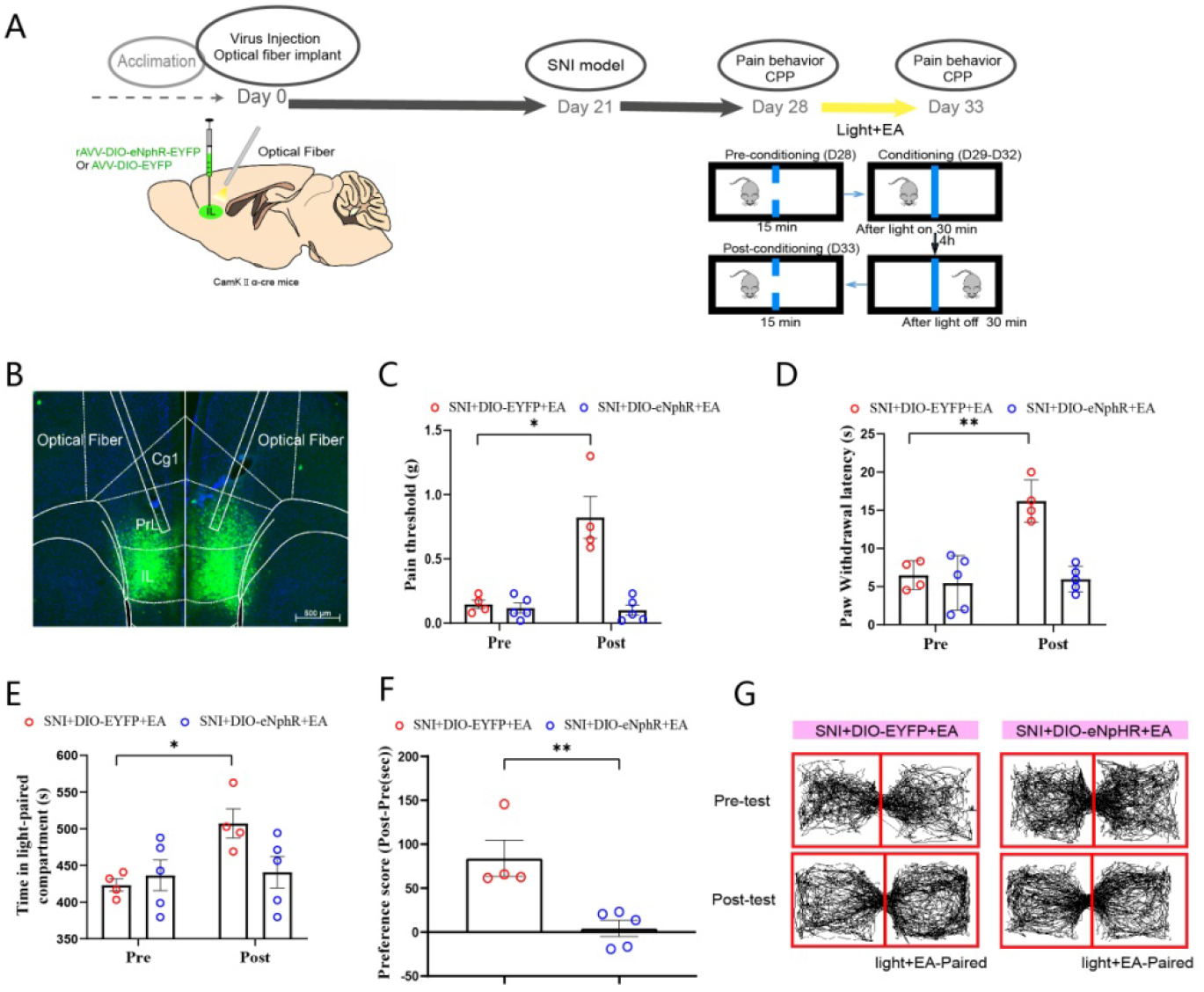
Inhibition of glutamatergic neurons in IL reversed the effects of EA in SNI mice. (A) Schematic diagram showing the timeline of experiment and the injection of AAV-EYFP or rAAV-eNpHR-EYFP into the bilateral IL and embedding of optical fibers in CamK II α-Cre mice. (B) Representative images of viral injection site and fiber embedding site in CamK II α-Cre mice. Bar = 500μm. (C)-(D) The increased thresholds of mechanical allodynia and thermal hyperalgesia induced by EA was inhibited after yellow light stimulation. (C) Pain threshold (g) before and after light stimulation. (D) Paw withdrawal latency (s) before and after light stimulation; n=4 mice in each group. (E)-(F) After four consecutive days of light stimulation, the increase of the residence time of EA-conditioned chamber were inhibited in SNI model mice. (E) Time in light-paired compartment (s) before and after light stimulation; (F) CPP scores. n=4 mice in each group. (G) The trajectory of SNI mice in a CPP apparatus during pre-test phase and post-test phase.*p < 0.05, **p < 0.01, ***p < 0.001. All data were presented as mean ± s.e.m. Two-way analyses of variance (ANOVAs) followed by Fisher’s LSD test and Unpaired Student’s t test were used for statistical analysis. All data were presented as mean ± s.e.m.

## Discussion

Peripheral neuromodulation elicited by EA can be considered as a highly specialized sensory experience. It can be precisely localized in somatotopy and described in modality and intensity [45]. Local sensory signals induced by EA recruit a basic sensory, stimulate peripheral nerve terminals, and carry processed information to a multitude of brain regions. However, whether signals of peripheral neuromodulation activating the brain’s neural system can translate into affective-motivational reactions is largely unknown. In this study, we systematically examined the affective-motivation responses of pain relief following application of EA stimulation in human and animals. We found that EA developed CPP based on relief from pain in different animal models of pain, which is consistent with clinical observations that the motivation and emotion could be induced by EA after pain relief in patients. These results provide proof of concept for the idea that peripheral neuromodulation signals reflect a multidimensional composite created by afferent sensory inputs and their processing to the brain circuitry, which evoked expectations, affective and motivational processing. Hence, EA is not only a sensory input but represents a dynamic state that shapes diverse psychosocial responses. These results may provide with multiple clinical implications, including treating pain conditions with psychiatric comorbidities, and understanding of EA-induced placebo effects.

EA-induced affective-motivational response of pain relief is clinically significant. Kong et al. suggest that conditioning positive expectation can amplify acupuncture analgesia as detected by subjective pain sensory rating changes [46]. Different from other studies [46,47], we assessed the effects of EA on affective and motivational response following EA treatment. After EA stimulation, patients with pain display profound affective and motivational responses (induced positive affective, enhanced motivation for EA, and reduced negative affect). The affective and motivational responses may evoke patients’ inherent power to alter the degree and quality of pain that is experienced and drive decisions to seek the next EA treatment. Furthermore, we found that EA still produced reinstatement of CPP even after the disappearance of pain in rats. This is the direct evidence in experimental animal studies that the persistent effects of EA on affective-motivational response may not require ongoing nociceptive signaling (independently from the perception of pain), which is consistent with our clinical observation that the patient still wanted to seek EA even after the disappearance of pain symptom. It should be noted that chronic pain can induce long term plasticity, substantially alter brain structural features, and also affect brain functional activity [31]. Our mechanism study focused on early stage of chronic pain. In fact, our previous data show that EA-treated rats exhibited marked preferences on days 7, 14, and 21 after SNI surgery (early stage), but not on days 28 and 45 after SNI surgery (late stage) [26], different from EA-induced antihyperalgesic effects. Hence, whether EA-induced affective-motivational response is correlated with the time window of chronic pain needs further research.

CPP has been induced by EA in different animal models of pain. These manipulations did not produce CPP in normal rats. It suggests that EA may become motivated in the presence of pain,but not EA is motivational intrinsically. This conclusion is consistent with the concept of negative reinforcement elicited by relief of an aversive state [47]. Also, the results of our behavioral experiments extend the conclusions of previous studies on the benign regulation of EA [48,49]. The variability in the response to EA might be explained partially by state-dependent effects of EA [49].

Neuromodulation alters nerve activity through targeted delivery of a stimulus to specific neurological sites. Brain imaging studies have demonstrated that EA can induce widespread brain activity changes. However, it is difficult to distinguish acupuncture-evoked widespread brain activation of the brain from the brain activity resulting from its therapeutic effect. In addition, the shortage of available neural circuit involving the effects of EA [2], coupled with a potentially broad use of EA, motivates the investigation we describe here. We identified glutamatergic neurons in the IL that underlies EA analgesia and affective-motivational responses. The IL plays pivotal roles in modulating goal-directed behaviors, emotional processes, and executive functions. IL activation also reflects a form of externally elicited top-down control that modulates sensory and affective processes of pain. Human and animal studies have shown that chronic pain induced decreased activity in the mPFC [17,18]. Stimulation of the mPFC produces descending pain control, including sensory and affective components [51]. Our lab has previously demonstrated that acupuncture can influence IL neuron firing activity [21,22]. Recent study demonstrates that the IL alters EA effect in animals with neuropathic chronic pain [52]. Herein, we additionally determined that activation of IL glutamatergic neurons mimicked EA-induced analgesia and conditioned place preference (CPP) behaviors, and inhibition of glutamatergic neurons in the IL reversed the effects of EA. Given that the clinical application of deep brain stimulation (DBS) are limited by the anesthetics and surgery, EA-induced peripheral neuromodulation may represent an alternative strategy for DBS as a means of activating the IL to treat symptoms associated with pain.

In summary, our data provide strong evidence that peripheral neuromodulation with EA induces analgesia and affective-motivational responses. EA-induced neuronal activity in the IL can drive EA analgesia and affective-motivational processes. These results illustrate an example in which emotional dimension of pain is directly influenced through the peripheral neuromodulation by EA, and provide the basis for EA that can precisely target the top-down neural control to relieve chronic pain in psychological and clinical situations.

## Data availability

GraphPad Prism and Matlab were used for all statistical analyses. The data that support the findings of this study are available on request from the corresponding author.

## Acknowledgments

This work was supported by Natural Science Foundation of China (81873379 and 82104990).

